# Temporal misexpression of *En1* during limb development causes distinct phenotypes

**DOI:** 10.1101/2024.08.06.606766

**Authors:** Alessa R. Ringel, Andreas Magg, Natalia Benetti, Robert Schöpflin, Mira Kühnlein, Asita Carola Stiege, Ute Fischer, Lars Wittler, Stephan Lorenz, Stefan Mundlos, Lila Allou

## Abstract

The precise spatiotemporal regulation of developmental genes is required for proper organogenesis. *Engrailed-1* (*En1*) is essential for dorsal-ventral patterning during mouse limb development from embryonic day E9.5 to E11.5. Previously, we identified the long non-coding RNA locus *Maenli*, which drives limb-specific *En1* expression at E9.5. In this study, we investigated the regulatory mechanisms sustaining *En1* expression at later developmental stages when *Maenli* transcriptional activity is drastically reduced. Using *in vivo* CRISPR editing, we identified two intergenic enhancer elements, LSEE1 and LSEE2, that maintain *En1* expression at E10.5 and E11.5. Mice lacking these enhancers exhibit only a subset of the limb malformations observed in *En1* and *Maenli* mutants, indicating that the timing of *En1* misexpression causes distinct phenotypes. These findings underscore the role of temporally restricted activities of *cis*-regulatory elements, including lncRNA loci and enhancers, in modulating gene expression and explaining subtle differences in complex disease phenotypes.

## Introduction

Healthy embryonic development relies on the precise expression of genes in space and time. Such spatiotemporal regulation of developmental genes is orchestrated by the action of *cis*-regulatory elements (CREs), including enhancers^1^ and long non-coding RNA (lncRNA) loci^2, 3^. Enhancers are short, non-coding DNA sequences which boost transcription rates from promoters and can be located at varying distances from their target genes^1, 4^. The interaction between enhancers and their target promoters is enabled by folding of the chromatin^5, 6^, and is often constrained to topologically associated domains (TADs). TADs are genomic regions that have higher chromatin interaction frequencies within themselves than the rest of the genome^7, 8^. In the recent years, lncRNA loci have gained recognition in regulating tissue-specific developmental gene expression^2, 3^. LncRNAs are noncoding transcripts longer than 500 bp that are typically expressed at low levels compared to mRNAs^2, 3^. Some lncRNA loci have been reported to function as CREs^2, 3^.

Developmental genes are often regulated by a complex interplay of multiple CREs that can cooperate in additive, synergistic, or redundant manners^9, 10^. Here, we deployed an *in vivo* CRISPR/Cas genome-engineering approach to dissect the regulatory architecture of the *Engrailed-1* (*En1*) gene during limb development. *En1* encodes the homeobox transcription factor EN1 which can act as both a transcriptional activator and a repressor^11^. It is involved in regionalization during early embryogenesis. As such, its precise spatiotemporal expression is crucial for the development of several tissues like bone, muscle, cartilage, and skin^11^. *En1* is first expressed at embryonic day (E) 8 in the mouse, in the midbrain/hindbrain junction. By E9.5, it is found in the mesenchyme adjacent to the midbrain/hindbrain junction, the developing pituitary gland, the somites, and the limb buds. In the limbs it is specifically expressed in the ventral ectoderm and apical ectodermal ridge^11^. It is also expressed in other structures (e.g. central nervous system, spinal cord, tail bud, mandibular arch etc.) from E10.5^11^.

Homozygous *En1* knockout mice die shortly after birth and display multiple developmental abnormalities, including brain and skeletal (sternum, rib and limb) defects^11, 12^. The limb phenotype manifests as polydactyly and syndactyly^11, 12^. Deleting only the homeobox-coding sequence of *En1* allows rare survival of mice which show a double dorsal limb phenotype; consisting of the dorsalization of ventral structures (ventral or circumferential claws, ventral hairs, loss of pads, ventral digits, hyperpigmentation), as well as poly– and syndactyly^13, 14^. These early reports described the defects as embryonic (syn– and polydactyly) and postnatal (double dorsal) limb phenotypes. However, as the embryonic phenotype is also present postnatally, in this study we refer to the poly-/syndactyly and double dorsal limb defects as *early* and *late* phenotypes respectively.

While the phenotypic consequences of *En1* loss in the embryonic limb have been well characterized^15^, less is known about the regulatory elements controlling *En1* expression. *En1* is positioned in a ∼ 650 kilobase (kb) TAD in which we previously identified the lncRNA locus *Maenli* (<u>M</u>aster <u>a</u>ctivator of *<u>e</u>ngrailed-1* in the <u>li</u>mb). *Maenli* initiates *En1* expression in the mouse limb bud at E9.5^15^. However, at E10.5 and E11.5, *Maenli* transcriptional activity significantly decreases, yet *En1* expression is maintained^15^, suggesting that additional regulatory elements may be present to sustain *En1* expression.

Here, we employed *in vivo* CRISPR/Cas9 genome editing to elucidate the regulatory elements responsible for maintaining *En1* expression. We have identified two enhancer elements which cooperate to drive *En1* expression in the E10.5 and E11.5 limb bud that we term <u>L</u>ate <u>S</u>tage *<u>E</u>ngrailed-1*-associated limb-specific <u>E</u>nhancers 1 and 2 (LSEE1 and LSEE2). Strikingly, simultaneous homozygous deletion of *LSEE1*&*2* causes morphological limb defects resembling only the late-stage *En1* mutant limb abnormalities^12^. In contrast, *Maenli*^−/−^ mice recapitulate both the early– and late-stage morphological limb defects, supporting a distinct spatiotemporal contribution of the CREs in controlling *En1* expression. Our findings show that LSEE1&2 play a crucial role in sustaining *En1* expression during limb development. This study highlights how CREs acting in a time-specific manner can contribute to genetic disorders with subtle phenotypic variations.

## Results

### Regulatory elements beyond *Maenli* contribute to *En1* expression in E10.5 and E11.5 limb buds

To investigate whether additional regulatory elements drive *En1* expression in E10.5 and E11.5 limb buds, we performed a genetic dissection of the *En1* regulatory landscape using *in vivo* CRISPR/Cas9 genome editing. We generated two large deletions (*Del1* and *Del2*) within the *En1* regulatory landscape, and a deletion of only the *Maenli* locus (*Del3*) **(Fig. 1a)**. *Del1* eliminates the genomic region between *En1* and *Maenli* while *Del2* spans from *Maenli* to the telomeric boundary of the *En1* TAD **(Fig. 1a)**.

**Figure 1.**
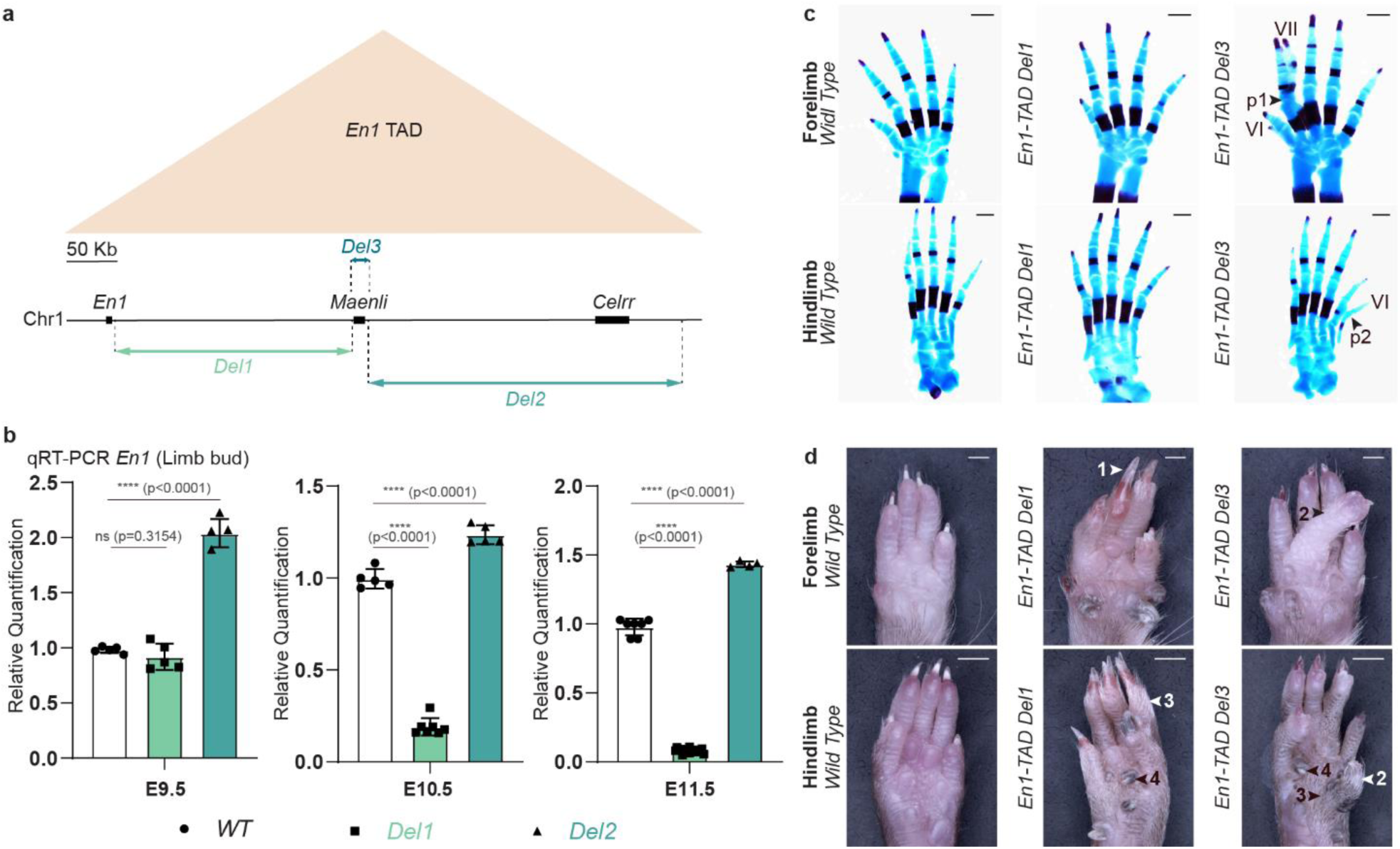
*Del1* significantly decreases *En1* expression in the limb and causes a double dorsal limb phenotype in mice. **a**, Schematic representation of the *En1* TAD (beige triangle) and the CRISPR-Cas9 genetic dissection of the *En1* regulatory landscape (*Del1*, *Del2*, and *Del3*). **b,** Normalized qRT-PCRs of *En1* in E9.5, E10.5, and E11.5 limb embryos show a significant loss of *En1* expression at E10.5 and E11.5 upon deletion of the genomic region between *En1* and *Maenli* (*Del1*). By contrast, this deletion did not significantly affect *En1* expression at E9.5. Deletion of the genomic region between *Maenli* and the telomeric end of the *En1* TAD (*Del2*) resulted in a significant increase of *En1* expression at E9.5, E10.5, and E11.5, respectively. Data were normalized to wild-type expression; one-tailed t-test; data are mean±SD; n = 5 *WT*, 5 *Del1*, 5 *Del2* at E9.5; n = 5 *WT*, 7 *Del1*, 5 *Del2* at E10.5; n = 7 *WT*, 18 *Del1*, 4 *Del2* at E11.5. ns, non-significant; *, (p < 0.01); p, p-value; WT, wild-type. **c,** Alcian blue (cartilage) and alizarin red (bone) stained limbs prepared from wild-type, *Del1*, and *Del3* E18.5 embryos. The *Del1* mutant is indistinguishable from the wild-type. The *Del3* mutant exhibits the presence of a polydactyly (digit VI) and an ectopic ventral digit (VII) fused at the level of phalange 1 in the forelimb, and an ectopic ventral digit (VI) fused at the level of phalange 2 in the hindlimb. p, phalange. Scale bars, 500 µm; n = 10 *WT*, 8 *Del1*, and 9 *Del3*. **d,** Ventral views of wild-type, *Del1*, and *Del3* adult (8-weeks) fore– and hindpaws illustrating the presence of ectopic ventral nails (1), ectopic ventral digits (2), and ectopic ventral hairs (3). The pigmented metatarsal pads (4) are elongated and hardened, resembling nails. Scale bars, 1,000 µm for forelimbs and 2,000 µm for hindlimbs, respectively; n = 13 *Del1* and 15 *Del3*.

In this study, qPCR analyses have been performed using *Rps9* as a housekeeping gene to normalize the expression levels of our target genes. We also used *Gapdh* as an endogenous control for all our qPCR measurements and observed no changes in *Gapdh* levels. These analyses showed that *Del1* and *Del2* did not affect *Maenli* expression in the E9.5 limb bud **(Extended Data Fig. 1a, b)**. However, although *Del1* did not significantly affect *En1* expression in the E9.5 limb bud, expression was significantly decreased at E10.5 and E11.5 **(Fig. 1** **b and Extended Data Fig. 1b, c)**. *Del2* caused a significant increase in *En1* expression at all three developmental stages **(Fig. 1** **b and Extended Data Fig. 1b, c)**.

Interestingly, *Del1* mice showed strong limb malformations resembling those previously described for *En1* mutant^12, 13^ and *Maenli^−/−^*^15^ mice. Notably, these mice exhibited the late-stage (∼ 4 weeks postnatal) morphological limb defects consisting of dorsalization of ventral structures, including ventral or circumferential claws, ventral hairs, and loss of pads. However, they did not exhibit the early-stage (< 0 days postnatal) morphological limb abnormalities, including syndactyly, polydactyly, and ventral digits **(Fig. 1c, d** **and Extended Data Fig. 1d, e)**. These data suggest that *Del1* harbours regulatory elements driving *En1* expression in the E10.5 and E11.5 limb bud, with their loss only causing the late-stage *En1* mutant limb phenotype.

### A second lncRNA locus, *Lmer,* is dispensable for limb-specific *En1* expression

To identify potential regulatory elements within *Del1* driving *En1* expression at E10.5 and E11.5, we first performed capture Hi-C (cHi-C) in E10.5 limb buds **(Fig. 2a)**. We examined chromatin interactions between *Del1* and *En1* using the *En1* promoter as viewpoint by deriving a virtual 4C profile from the cHi-C data. However, despite strong interactions between the *En1* promoter with the telomeric end of the *En1* TAD we found no prominent specific interaction with a particular region in *Del1* **(Fig. 2a)**.

**Figure 2.**
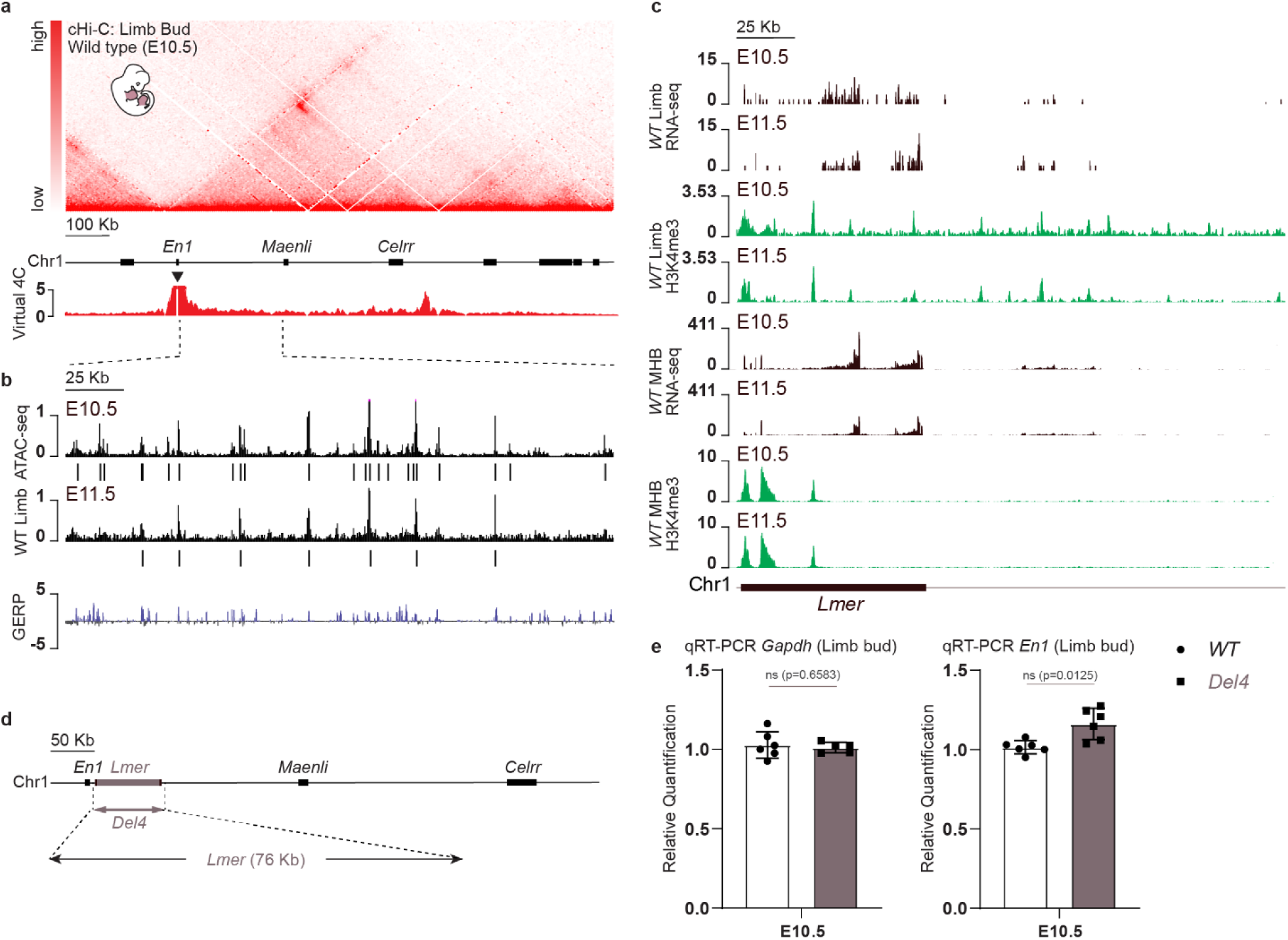
*Lmer* is dispensable for *En1* expression regulation during limb development. **a**, cHi-C map of the mouse limb buds at E10.5 shows a large TAD encompassing the *En1* gene and the *Maenli* and *Celrr* lncRNA loci. Virtual 4C profile using the *En1* promoter as viewpoint (black triangle) shows no prominent interaction between *En1* and the genomic region upstream of *Maenli*. n = 2 biologically independent wild-type replicates with 2 technical replicates each. **b,** ATAC reads from wild-type (WT) E10.5 and E11.5 mouse limb buds are shown. Peak calling identifying genomic regions which are enriched in aligned ATAC reads (to a statistically significant extent) compared to background levels is also shown. n = 2 biologically independent wild-type replicates. **c,** Poly(A)^+^ RNA-seq and H3K4me3 ChIP-seq profiles of E10.5 and E11.5 mouse limb buds and midbrain-hindbrain junction show the presence of the lncRNA locus *Lmer*. n = 2 biologically independent wild-type replicates. **d,** Schematic representation of the CRISPR-Cas9 genetic deletion involving the lncRNA locus *Lmer* (*Del4*). **e,** Normalized qRT-PCRs of *Gapdh* and *En1* in E10.5 mouse limb embryos show no significant changes in *Gapdh* and *En1* expression upon deleting the lncRNA locus *Lmer* (*Del4*). Data were normalized to wild-type expression; one-tailed t-test; data are mean±SD; n = 6 *WT*, 6 *Del4* at E10.5 for *En1* expression quantification; n = 6 *WT*, 5 *Del4* for *Gapdh* expression quantification. ns, non-significant; p, p-value; WT, wild-type.

We then searched for putative enhancers and transcripts in *Del1* by analysing previously published^15, 16^ assay for transposase-accessible chromatin, followed by sequencing (ATAC-seq) data in E10.5 and E11.5 limb buds. *Del1* was found to contain 7-23 putative enhancers as defined by ATAC-seq peaks **(Fig. 2b)**. Additionally, we performed poly(A)^+^ RNA sequencing (RNA-seq) in wildtype E10.5 and E11.5 limb buds, somites, and the midbrain/hindbrain junction. These analyses revealed the presence of a transcript in the limb buds and the midbrain/hindbrain junction within *Del1* that was not detected in the somites **(Fig. 2c** **and Extended Data Fig. 2a)**. PhyloCSF analysis showed that this transcript lacks protein-coding potential, suggesting that it is a lncRNA **(Extended Data Fig. 2b)**. The low conservation of its nucleotide sequence with only 28% similarity between mice and humans and low conservation across other species supports this **(Extended Data Fig. 2b)**. We named this lncRNA locus *Lmer* (<u>L</u>imb <u>m</u>idbrain/hindbrain junction <u>e</u>xpressed <u>R</u>NA) **(Fig. 2c)**.

The mouse *Lmer* locus encompasses 11 exons spanning approximately 76 kb and is located around 6 kb downstream of the 3’-end of *En1* **(Fig. 2d)**. Its transcriptional start site (TSS) overlaps with H3K4me3 marks in E10.5 and E11.5 limb buds and the midbrain/hindbrain junction **(Fig. 2c)**. We tested if the *Lmer* locus is required for *En1* expression and limb development *in vivo* by deleting the *Lmer* locus itself (*Del4*) **(Fig. 2d)**. This deletion did not significantly affect *En1* expression in E10.5 limb buds **(Fig. 2e)**. Thus, the mouse *Lmer* locus is dispensable for the correct timing and location of *En1* expression during limb development.

### Subtle differences in limb defects reveal a critical region for regulating late-stage *En1* expression

We next hypothesized that the genomic region between *Lmer* and *Maenli* (∼ 157 kb) contains regulatory elements controlling *En1* expression in E10.5 and E11.5 limb buds. This genomic region contains 4-13 putative enhancers in the limbs as defined by ATAC-seq peaks **(Fig. 2b)**. To determine the position of potential *En1* regulatory elements within this large genomic region, we analysed the phenotypic differences between three mutants (*Del1* and *Del3* generated in this study, and *Del27* generated in our previous study^15^) **(Extended Data Fig. 3a)**. Interestingly, while all of these mutants exhibited limb abnormalities resembling those of *En1* mutant mice^12, 13^ (complete penetrance), they showed small but noticeable phenotypic differences.

*Del27* mice (with a deletion of the *Maenli* locus and ∼ 9 kb of the adjacent noncoding DNA sequences) exhibited syndactyly with or without polydactyly (early-stage limb defects) in all forelimbs and ectopic ventral nails (late-stage limb abnormalities) in all forelimb and hindlimb digits (n = 16)^15^. In *Del3* mutants, however, the syndactyly with or without polydactyly was observed in ∼ 1 forelimb of each embryo and ectopic ventral nails in ∼ 2-3 digits of each limb (n = 15). Finally, in *Del1* mice, the syndactyly and polydactyly were absent, with ectopic ventral nails present in all forelimb and hindlimb digits (n = 13) **(Extended Data Fig. 3a)**. These observations suggest temporal differences in *En1* expression between the three mutants during limb development.

We tested this by quantifying *En1* expression at E9.5, E10.5, and E11.5 in *Del3* and *Del27* limb buds using RNA-seq and *Del3* limb mutants using qRT-PCR **(Extended Data Fig. 3b, c)**. *Del3* embryonic limbs showed an almost complete loss of *En1* expression at E9.5 and reduced levels at E10.5 and E11.5 with ∼ 35% and ∼ 62% of wildtype respectively **(Extended Data Fig. 3b, c)**. Moreover, RNA-seq in *Del27* limb buds showed a complete loss of *En1* expression at all three developmental stages **(Extended Data Fig. 3c)**. These data demonstrate that *Del27* contains the complete regulatory information required to control *En1* expression during limb development. In contrast, deletion of *Maenli* alone (*Del3*) leads to only a partial loss of *En1* expression in E10.5 and E11.5 limb buds, confirming that additional regulatory elements maintain *En1* expression. In combination with the results from the *Del1* mutants **(Fig. 1** **b and Extended Data Fig. 1b, c)**, these findings suggest that putative regulatory elements located within the overlapping genomic region between *Del1* and *Del27* control *En1* expression in the limb buds after E9.5.

### Late-stage enhancers are required for proper dorsal-ventral limb patterning

The overlapping 6 kb genomic region between *Del1* and *Del27* contains one putative enhancer element as defined by ATAC-seq **(Fig. 3a)**. A ∼ 2 kb deletion of this putative enhancer (*Del5*) led to a significant decrease in *En1* expression by ∼ 23% in E10.5 mutant limb buds **(Fig. 3a, b)**. This reduction was less severe than the one observed in E10.5 *Del1* limb mutants **(Fig. 1b** **and Extended Data Fig. 1b, c)**. Therefore, we next generated a ∼ 4 kb deletion covering both conserved noncoding DNA elements as defined by vertebrate conservation measured using Genomic Evolutionary Rate Profiling (GERP) (*Del6*) **(Fig. 3a)**. This significantly decreased *En1* expression in E10.5 and E11.5 limb buds (∼ 77% and ∼ 91% reduction respectively) **(Fig. 3c, d)**. *Del6* mutants displayed the same reduction in *En1* expression in E10.5 and E11.5 limb buds as *Del1* mutants, and also only displayed the late-stage limb malformations (n = 7) **(Fig. 1b, d, 3c, d, e and Extended Data Fig. 1b, c, e)**. These data indicate that the deleted region (*Del6)* contains enhancers which cooperate to drive *En1* expression in the E10.5 and E11.5 limb buds. We named these enhancer elements LSEE1 and LSEE2 (<u>L</u>ate <u>S</u>tage *<u>E</u>ngrailed-1*-associated limb-specific <u>E</u>nhancer) respectively **(Fig. 3a)**. Given that LSEE1&2 are positioned ∼ 2.5 kb upstream of *Maenli*, we next examined whether the initial transcriptional activity of *Maenli* at E9.5 is needed for the downstream activity of LSEE1&2. We generated a ∼ 235 kb inversion (*Inv*) relocating the enhancers from the 3’ end of *Maenli* to the 3’ end of *En1* (∼4 kb downstream) **(Extended Data Fig. 4a)**. This inversion had no significant impact on *En1* expression in E11.5 limb buds **(Extended Data Fig. 4b)**, suggesting that LSEE1&2 enhancer activity may not depend on the initial transcriptional elongation through the *Maenli* locus.

**Figure 3.**
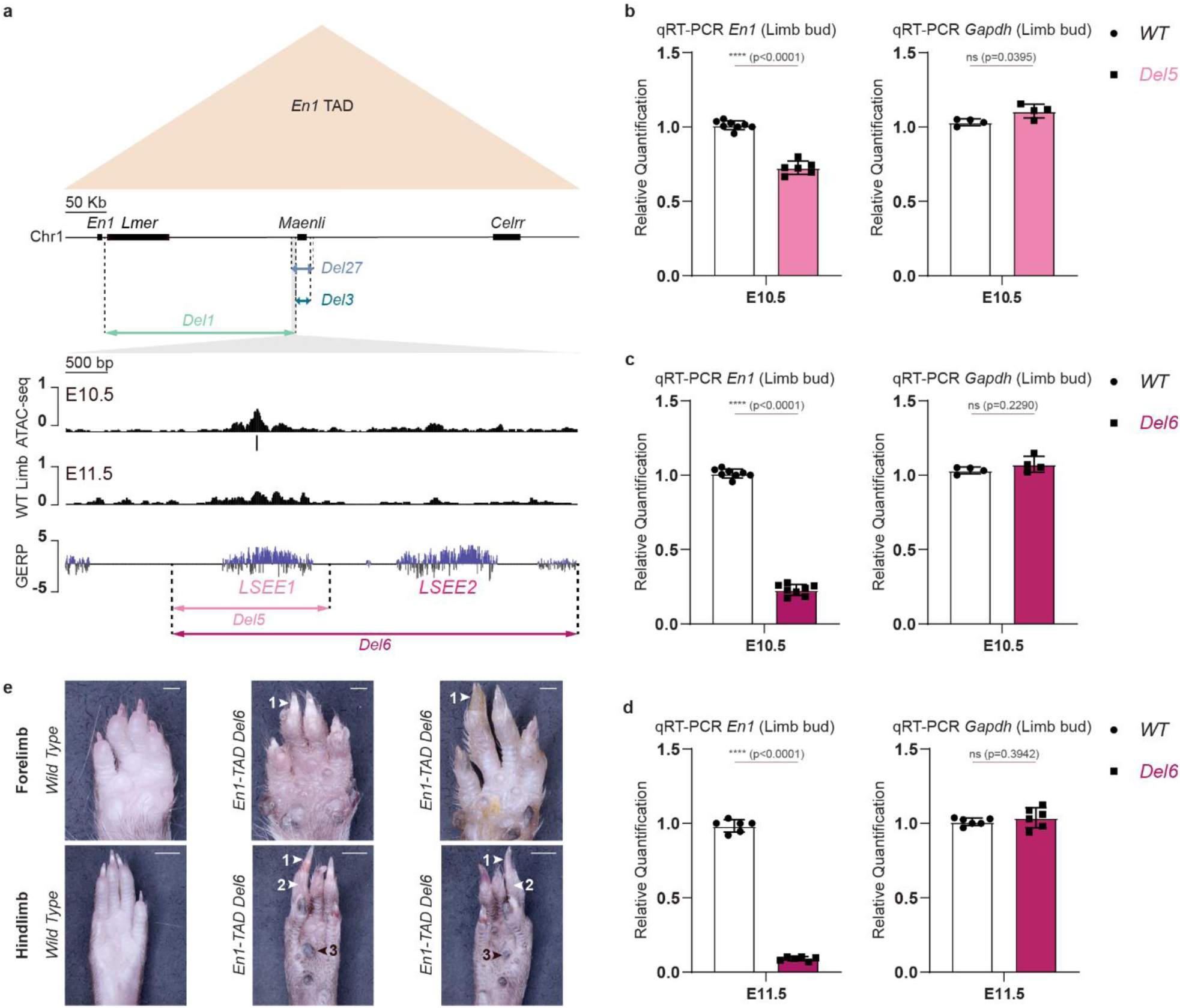
LSEE1 and LSEE2 enhancers cooperate to drive *En1* expression in E10.5 and E11.5 mouse limb buds. **a**, Schematic representation of the *En1*-TAD (beige triangle) and the CRISPR-Cas9 genetic deletions (*Del1*, *Del3*, and *Del27*) affecting *En1* limb expression and causing abnormal limb bud dorsoventral patterning in mice. The overlapping genomic region between *Del1* and *Del27* is highlighted in grey, and a zoom-in of the region is shown. ATAC reads from wild-type E10.5 and E11.5 mouse limb buds indicate the presence of a putative enhancer element within the overlapping genomic region. Vertebrate conservation is measured using the Genomic Evolutionary Rate Profiling (GERP) method. It indicates the presence of two highly conserved non-coding DNA elements within the overlapping genomic region. Finally, a schematic representation of the CRISPR-Cas9 genetic dissection of the overlapping genomic region (*Del5* and *Del6*) is shown. **b,** Normalized qRT-PCRs of *En1* and *Gapdh* in E10.5 mouse limb embryos show no significant changes in *Gapdh* expression, but a significant loss of *En1* expression upon deleting the single enhancer LSEE1 (*Del5*). Data were normalized to wild-type expression; one-tailed t-test; data are mean±SD; n = 8 *WT*, 6 *Del5* for *En1* expression quantification; n = 4 *WT*, 4 *Del5* for *Gapdh* expression quantification. ns, non-significant; *, (p < 0.01); p, p-value; WT, wild-type. **c, d,** Normalized qRT-PCRs of *En1* and *Gapdh* in E10.5 and E11.5 mouse limb buds isolated from *Del6* embryos show no significant changes in *Gapdh* expression, but a significant loss of *En1* expression at both developmental stages with an almost complete loss at E11.5. Data were normalized to wild-type expression; one-tailed t-test; data are mean±SD; (n = 8 *WT*, 8 *Del6* at E10.5; n = 6 *WT*, 6 *Del6* at E11.5 for *En1* expression quantification); (n = 4 *WT*, 4 *Del6* at E10.5; n = 6 *WT*, 6 *Del6* at E11.5 for *Gapdh* expression quantification). ns, non-significant; *, (p < 0.01); p, p-value; WT, wild-type. **e,** Ventral views of wild-type and *Del6* adult (6-weeks) fore– and hind-paws illustrating the presence of ectopic ventral nails (1) and ectopic ventral hairs (2). The pigmented metatarsal pads (3) are elongated and hardened, resembling nails. Scale bars, 1,000 µm for forelimbs and 2,000 µm for hindlimbs, respectively; n = 7 *Del6*.

## Discussion

We previously characterized the lncRNA locus *Maenli* as the master activator of *En1* expression in the mouse limb bud at E9.5. Here, we show that *En1* expression is maintained at later developmental stages by two enhancers, LSEE1 and LSEE2. Overall, we show that *En1* expression during limb development is regulated by the coordinated action of both a lncRNA locus and enhancers over time. To our knowledge, this is the first example of time-specific gene regulation within the same tissue being linked to distinct congenital malformations.

Deleting the genomic region between *Maenli* and the telomeric end of the *En1* TAD (*Del2*) significantly increased *En1* expression during limb development **(Fig. 1b** **and Extended Data Fig. 1b, c)**. This deletion shifts the telomeric TAD boundary closer to CREs of *En1,* possibly increasing their contact frequency with the *En1* promoter, leading to its upregulation. By contrast, removal of the region between *En1* and *Maenli* (*Del1*) significantly reduced *En1* expression in the E10.5 and E11.5 limb bud **(Fig. 1b** **and Extended Data Fig. 1b, c)**. In this region we found another lncRNA locus which we thought may regulate *En1* at later stages **(Fig. 2c)**. However, we found this lncRNA locus, *Lmer,* to be dispensable for limb development (*Del4*) **(Fig. 2d, e)**. Instead, *Lmer* is highly expressed in the midbrain/hindbrain junction, where *En1* is also expressed and is essential for its development^11^ **(Fig. 2c)**. How *En1* expression is controlled in this linear structure between the two brain regions is unknown. Hence, it is tempting to speculate that *Lmer* controls *En1* expression in the midbrain/hindbrain junction.

With further deletions we identified a ∼ 6 kb genomic region containing the LSEE1&2 enhancers required for maintaining *En1* expression after E9.5 **(Fig. 3a, b, c, d** **and Extended Data Fig. 3a, b, c)**. Homozygous deletion of LSEE1&2 resulted in up to 91% loss of *En1* expression in E11.5 limb buds **(Fig. 3d)**. Although we do not know whether LSEE1&2 cooperate redundantly, additively, or synergistically, it seems that their function is reliant on *Maenli* transcriptional activity. Deletion of *Maenli* and LSEE1&2 together (*Del27*) fully ablates *En1* expression in the limb **(Extended Data Fig. 3c)**. However, the presence of only LSEE1&2 in *Del3* (homozygous deletion of *Maenli*) causes a ∼92% reduction in *En1* expression at E9.5, but recovers to only ∼62% by E11.5 **(Extended Data Fig. 3b, c)**. This suggests that *En1* at later stages is driven by a combination of the enhancers and residual *Maenli* activity^15^ and that *Maenli* activity itself is required for the activity of the downstream enhancers.

A complex regulatory architecture with inter-dependencies on multiple *cis* regulatory elements *in vivo* has been reported only for few genes including *Sox9*^17^. *Sox9* plays a crucial role in sex determination where its transcription in the gonads relies on an enhancer cascade involving the activity of an early-enhancer that is required for later enhancer activation^17^. Here we also see a temporal interplay of CRE activity at the *En1* locus where the activity of *Maenli* is required for later enhancer activity. Based on the following two lines of reasoning, we hypothesize a transcription-based mechanism underlies LSEE1&2 activation. Firstly, we previously showed that insertion of a polyadenylation termination sequence shortly after the *Maenli* transcription start site ablated *En1* expression and H3K4me3 deposition across the *Del1* region^15^. However, inversion of this polyadenylation termination sequence did not affect *En1* expression levels. Although these experiments cannot distinguish between the act of *Maenli* transcription and the function of the transcript itself, they suggest that a transcriptional mechanism underlies *En1* activation and the establishment of a permissive chromatin state. Secondly, in this study we show that inverting the *Del1* region to bring LSEE1&2 ∼300kb away from *Maenli* had no effect on *En1* expression **(Extended Data Fig. 4a, b)**, indicating that *Maenli* transcription *per se* is not needed for LSEE1&2 activation. However, since this inversion placed LSEE1&2 ∼4kb downstream of *En1* **(Extended Data Fig. 4a)**, transcriptional elongation originating from *En1* may be sufficient for their activation. Thus, this experiment does not rule out a transcription-based mechanism of enhancer activation.

Another hypothesis is that *Maenli*’s promoter could be a *cis*-acting transcriptional stabilizer of *En1*. In this scenario, close physical proximity of the *Maenli* promotor with the *En1* promoter would also bring LSEE1&2 close to *En1*, therefore allowing them to sustain *En1* expression. Indeed, *Maenli* and *En1* are in close proximity with one another in the E9.5 limb bud^15^. This type of regulatory mechanism has been reported for the lncRNA locus *HASTER* which drives the interaction of enhancers with its adjacent target gene, *HNF1a*^18^. Alternatively, *En1* expression could be maintained during limb development through a positive feedback loop where EN1 binds to its own enhancers LSEE1&2. Residual *En1* expression (∼8%) in the absence of *Maenli* (*Del3*) could be sufficient to facilitate a low level of enhancer activity which would explain the partial recovery of *En1* expression at later stages **(Extended Data Fig. 3b, c)**. Such a positive feedback mechanism has been reported for LMX1B where it binds to two conserved *Lmx1b*-associated CREs which in turn amplify *Lmx1b* expression during limb dorsalization^19^. A more complex reinforcing feedback loop is also possible, where EN1 activates other transcription factors involved in ventral ectoderm and apical ectodermal ridge specification. Such a mechanism has been reported for *Fgf8* and *Fgf10* during limb development^20^. FGF10 signaling triggers the expression of *Fgf8* in the ectoderm through a positive regulatory loop, possibly coordinated via gene regulatory components.

*En1* mutation experiments from the 90s revealed distinct early (syndactyly, polydactyly) and late (double-dorsal) limb phenotypes in mice^14, 21^. Our previously reported mutant mice that lack *Maenli* and the here identified enhancers LSEE1&2 (*Del27*) displayed a syndactyly with or without polydactyly (early-stage limb defects) in all forelimbs, and ectopic ventral nails (late-stage limb abnormalities) in all forelimb and hindlimb digits^15^. However, in *Del3* mice (with only the *Maenli* deletion), the syndactyly with or without polydactyly was observed in ∼ 1 forelimb of each embryo and ectopic ventral nails in ∼ 2-3 digits of each limb **(Fig. 1c, d** **and Extended Data Fig. 1d, e)**, suggesting that LSEE1&2 partially rescue *En1* expression in the absence of *Maenli*. Interestingly, mice lacking LSEE1&2 (*Del1* and *Del6*) only display the late-stage morphological phenotype (**Fig. 1c, d, 3e** **and Extended Data Fig. 1d, e)**, suggesting that *En1*’s role at E9.5 is not affected, when establishment of proper digit number and separation occurs. Such a temporally restricted function of *En1* has also been suggested by homozygous replacement of *En1* with *En2*, which causes only the late-stage morphological phenotype^14, 21^. This suggests that the biochemical function of EN2 is able to compensate for that of EN1 for patterning proper digit number and separation, but not dorsal-ventral patterning. Here we show that these distinct temporal roles of *En1* are controlled by the coordinated function of *Maenli* and LSEE1&2 **(Fig. 4)**. However, to uncover the precise molecular mechanism(s) of *Maenli* and LSEE1&2 enhancer function and interplay further studies are necessary. One major limiting factor to these studies is the restriction of *En1* expression to a small number of cells in the limb bud (ventral ectoderm and apical ectodermal ridge) and the low level of *Maenli* expression.

**Figure 4.**
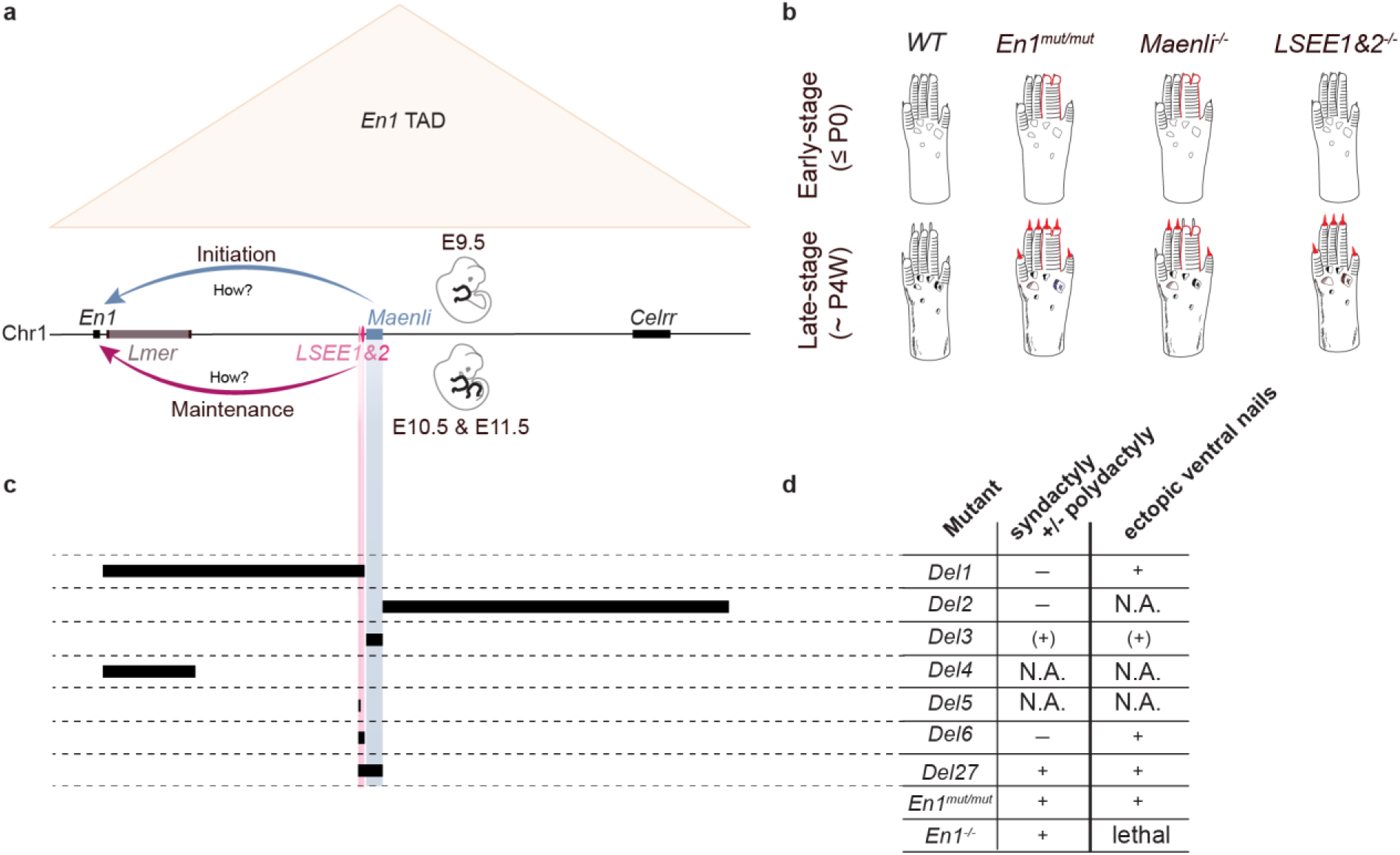
A model for limb-specific *En1* expression regulation. **a**, During limb development, *Maenli*, LSEE1, and LSEE2 fine-tune temporal *En1* expression. The lncRNA locus *Maenli* licenses *En1* expression specifically at E9.5. LSEE1&2 enhancers cooperate to drive *En1* expression, specifically at E10.5 and E11.5. The molecular mechanisms of *En1* expression control by *Maenli* and LSEE1&2 remain unknown (?). **b,** Schematic representation of *Maenli*^−/−^ and *LSEE1&2*^−/−^-associated limb phenotypes. *Maenli*^−/−^ and *LSEE1&2*^−/−^ mice exhibit limb defects recalling those of *En1* mutant mice. However, while *Maenli*^−/−^ mutants exhibit early– and late-stage morphological limb defects, *LSEE1&2*^−/−^ mutants exhibit only late-stage morphological limb abnormalities. Differential phenotypic features between *En1* mutant, *Maenli*^−/−^, and *LSEE1&2*^−/−^ mice include the syndactyly with or without polydactyly and ectopic ventral nails (highlighted in red). P0, postnatal day 0; P4W, 4 weeks postnatally. **c,** Schematic representation of the CRISPR-Cas9 genetic deletions generated in this study and previously (*Del27*)^15^. **d,** Table summarizing and showing differences in early– and late-stage phenotypes according to the different mutations. The phenotypes of *Del27*, *En1*^mut/mut^, and *En1*^−/−^ mice have been described in previous studies^12, 13, 15^. +, a syndactyly with or without polydactyly was observed in all forelimbs and ectopic ventral nails in all forelimb and hindlimb digits; (+), a syndactyly with or without polydactyly was observed in ∼ 1 forelimb of each embryo and ectopic ventral nails in ∼ 2-3 digits of each limb; –, the syndactyly and polydactyly were absent in all forelimbs; N.A., not assessed.

Overall, our findings have important implications for congenital limb malformations which are common^22^ and frequently isolated. The diagnostic yield in these cases remains low due to, at least in part, a focus on studying coding sequences. Our previous findings and others have shown that tissue-specific CREs are potential candidates to explain these isolated limb malformations^15, 19, 23, 24, 25, 26^. The findings of this study support the notion that time-specific CREs, including lncRNA loci and enhancers, are potential candidates for explaining isolated limb malformations with subtle phenotypic differences. Here we extend previous reports^18, 27, 28, 29^ by describing the coordinated action of a lncRNA locus and enhancers to control temporal gene expression. To our knowledge, this is the first study linking this temporal gene regulation to distinct phenotypes.

**Extended Data Figure 1.**
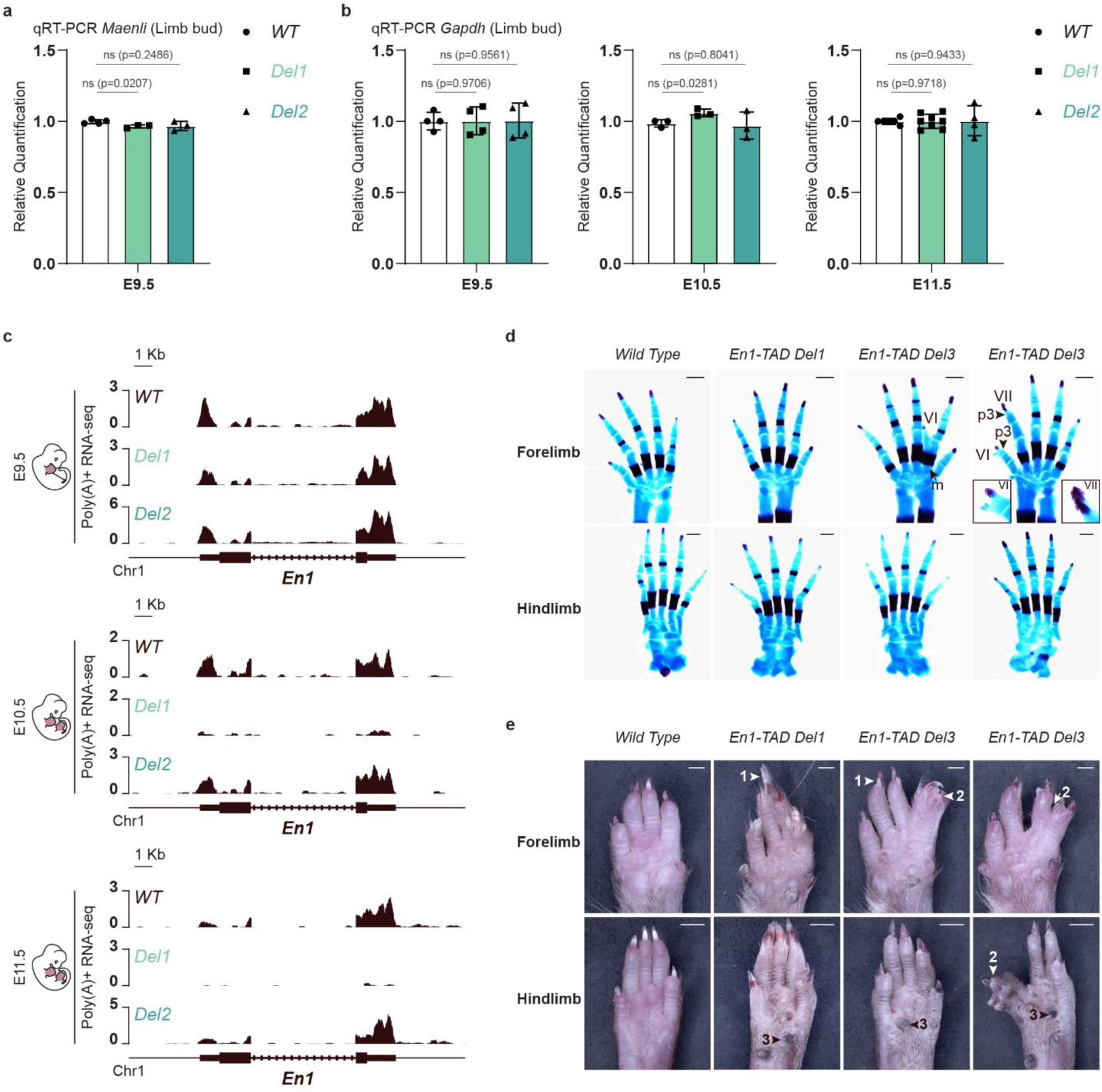
*Del1* mouse model exhibits abnormal limb bud dorso-ventral patterning similar to *Maenli*^−/−^ and *En1* mutant mice. **a**, Normalized qRT-PCRs of *Maenli* in E9.5 mouse limb embryos show that *Del1* and *Del2* did not significantly affect *Maenli* expression. Data were normalized to wild-type expression; one-tailed t-test; data are mean±SD; n = 4 *WT*, 3 *Del1*, 3 *Del2* at E9.5. ns, non-significant; p, p-value; WT, wild-type. **b,** Normalized qRT-PCRs of *Gapdh* in E9.5, E10.5, and E11.5 mouse limb embryos show no significant changes in *Gapdh* expression upon *Del1* and *Del2* deletions. Data were normalized to wild-type expression; one-tailed t-test; data are mean±SD; n= 4 *WT*, 4 *Del1*, 4 *Del2* at E9.5; n= 3 *WT*, 3 *Del1*, 3 *Del2* at E10.5; n = 6 *WT*, 8 *Del1*, 4 *Del2* at E11.5. ns, non-significant; p, p-value; WT, wild-type. **c,** Poly(A)^+^ RNA-seq profiles of E9.5, E10.5, and E11.5 mouse limb buds show that *Del1* did not affect *En1* expression at E9.5 but resulted in an almost complete loss of *En1* expression at E10.5 and E11.5. By contrast, *Del2* increased *En1* expression at E9.5, E10.5, and E11.5. WT, wild-type. n = 2 biologically independent *WT*, *Del1*, and *Del2* replicates. **d,** Alcian blue (cartilage) and alizarin red (bone) stained limbs prepared from wild-type, *Del1*, and *Del3* E18.5 embryos. The *Del1* mutant is indistinguishable from the wild-type. The *Del3* mutant exhibits the presence of ectopic ventral digits (VI and VII) fused at the level of metacarpal or phalanges 3 in the forelimb. m, metacarpal; p, phalange. Scale bars, 500 µm; n = 10 *WT*, 8 *Del1*, and 9 *Del3*. **e,** Ventral views of wild-type, *Del1*, and *Del3* adult (8-weeks) fore– and hind-paws illustrating the presence of ectopic ventral nails that manifest as circumferential nails or ventral nails formed opposite to the dorsal nails (1), and syndactyly (2). The pigmented metatarsal pads (3) are elongated and hardened, resembling nails. Scale bars, 1,000 µm for forelimbs and 2,000 µm for hindlimbs, respectively; n = 13 *Del1* and 15 *Del3*.

**Extended Data Figure 2.**
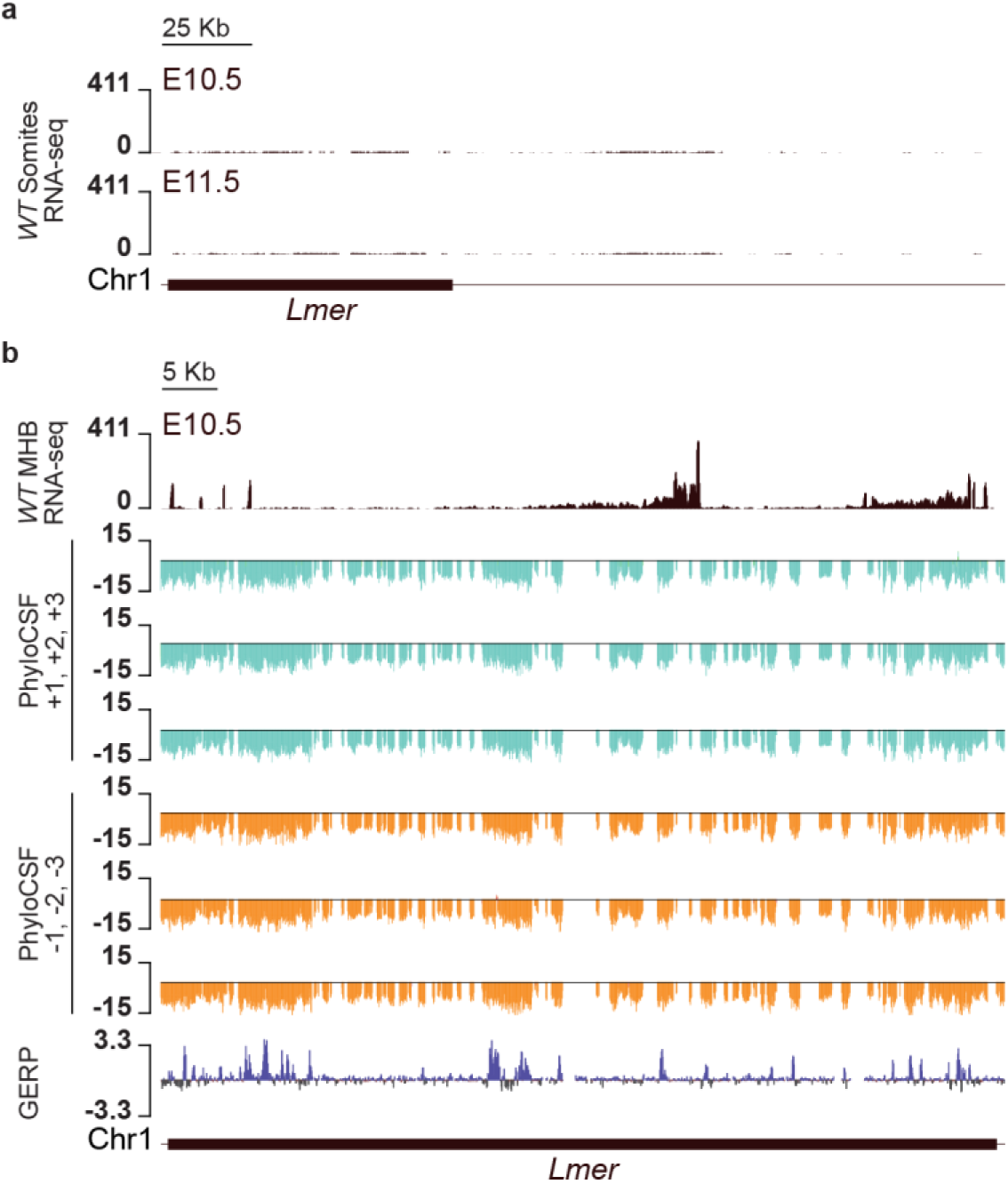
*Lmer* is expressed in the limb and midbrain-hindbrain boundary but not in the somites. **a**, Poly(A)^+^ RNA-seq profiles of E10.5 and E11.5 wild-type (WT) mouse somites show no expression of *Lmer* in these embryonic structures. n = 2 biologically independent wild-type replicates. **b,** Poly(A)^+^ RNA-seq profile of E10.5 wild-type (WT) mouse midbrain-hindbrain junction show a Zoom in of the *Lmer* transcript (exon-intron composition). PhyloCSF tracks show a lack of protein coding potential for the *Lmer* transcript. Vertebrate sequence conservation tracks measured using the Genomic Evolutionary Rate Profiling (GERP) method show that the *Lmer* nucleotide sequence (exon composition) exhibits little conservation. n = 2 biologically independent wild-type poly(A)^+^ RNA-seq replicates.

**Extended Data Figure 3.**
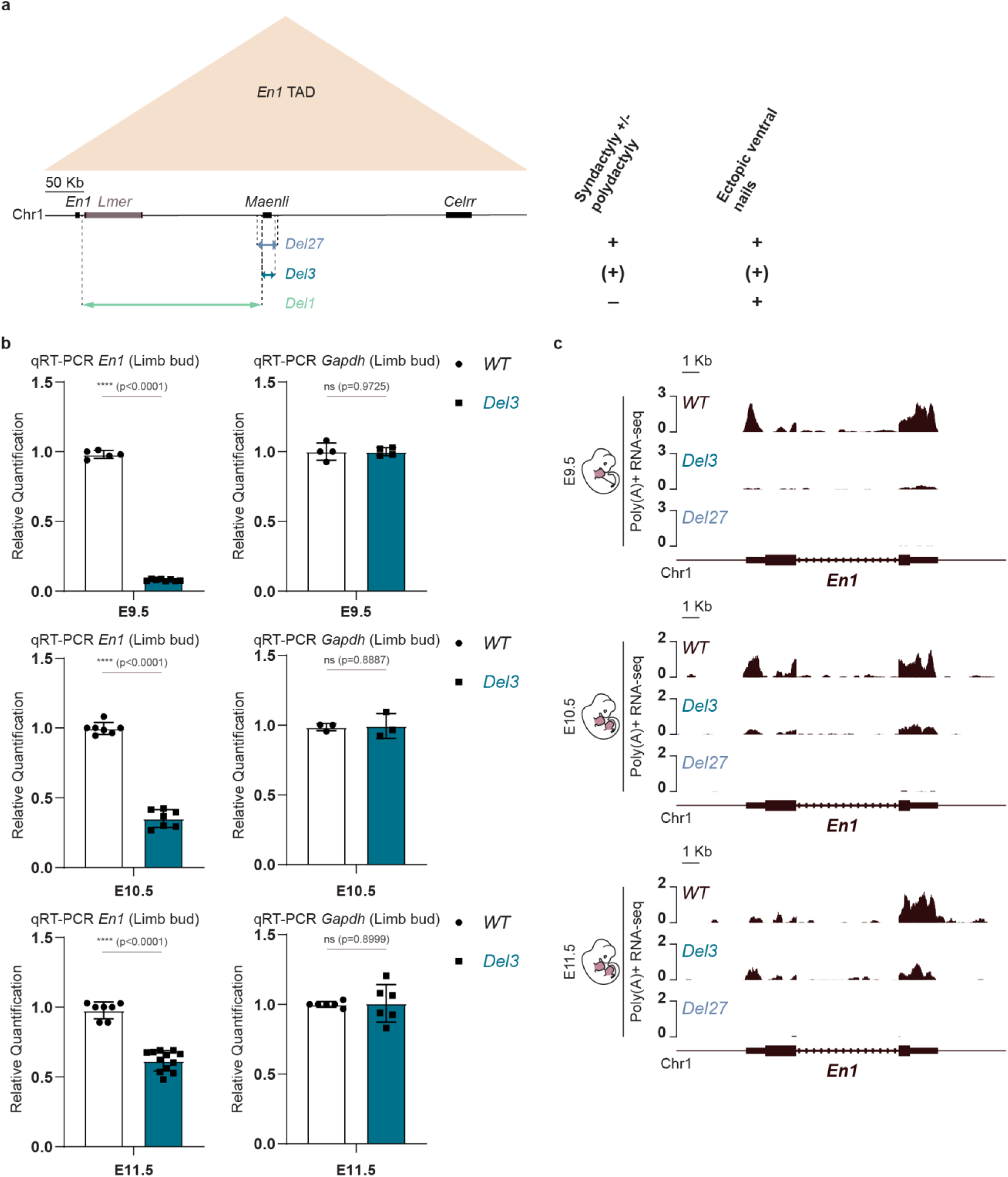
The overlapping genomic region between *Del1* and *Del27* contains the putative enhancers controlling *En1* expression in E10.5 and E11.5 mouse limb buds. **a**, Schematic representation of the *En1* TAD (beige triangle) and the CRISPR-Cas9 genetic deletions affecting *En1* limb expression and causing abnormal limb bud dorsoventral patterning in mice. The main differential phenotypic features of the mutants at E18.5 (syndactyly with or without polydactyly) and 8 weeks of age (ectopic ventral nails) are shown on the right. +, a syndactyly with or without polydactyly was observed in all forelimbs and ectopic ventral nails in all forelimb and hindlimb digits; (+), a syndactyly with or without polydactyly was observed in ∼ 1 forelimb of each embryo and ectopic ventral nails in ∼ 2-3 digits of each limb; –, the syndactyly and polydactyly were absent in all forelimbs. **b,** Normalized qRT-PCRs of *En1* and *Gapdh* in E9.5, E10.5, and E11.5 mouse limb embryos show no significant changes in *Gapdh* expression upon deleting *Maenli* (*Del3*). By contrast, this deletion significantly affected *En1* expression at E9.5, E10.5, and E11.5, with an almost complete loss at E9.5. *En1* expression was partially rescued at E10.5 and E11.5. Data were normalized to wild-type expression; one-tailed t-test; data are mean±SD; (n = 5 *WT*, 8 *Del3* at E9.5; n = 7 *WT*, 7 *Del3* at E10.5; n = 7 *WT*, 12 *Del3* at E11.5 for *En1* expression quantification); (n = 4 *WT*, 4 *Del3* at E9.5; n = 3 *WT*, 3 *Del3* at E10.5; n = 6 *WT*, 6 *Del3* at E11.5 for *Gapdh* expression quantification). ns, non-significant; *, (p < 0.01); p, p-value; WT, wild-type. **c,** Poly(A)^+^ RNA-seq profiles of E9.5, E10.5, and E11.5 mouse limb buds show that *Del27* caused a complete loss of *En1* expression at all three developmental stages. *Del3* caused an almost complete loss of *En1* expression at E9.5 with a partial rescue at E10.5 and E11.5. WT, wild-type. n = 2 biologically independent *WT*, *Del3*, and *Del27* replicates.

**Extended Data Figure 4.**
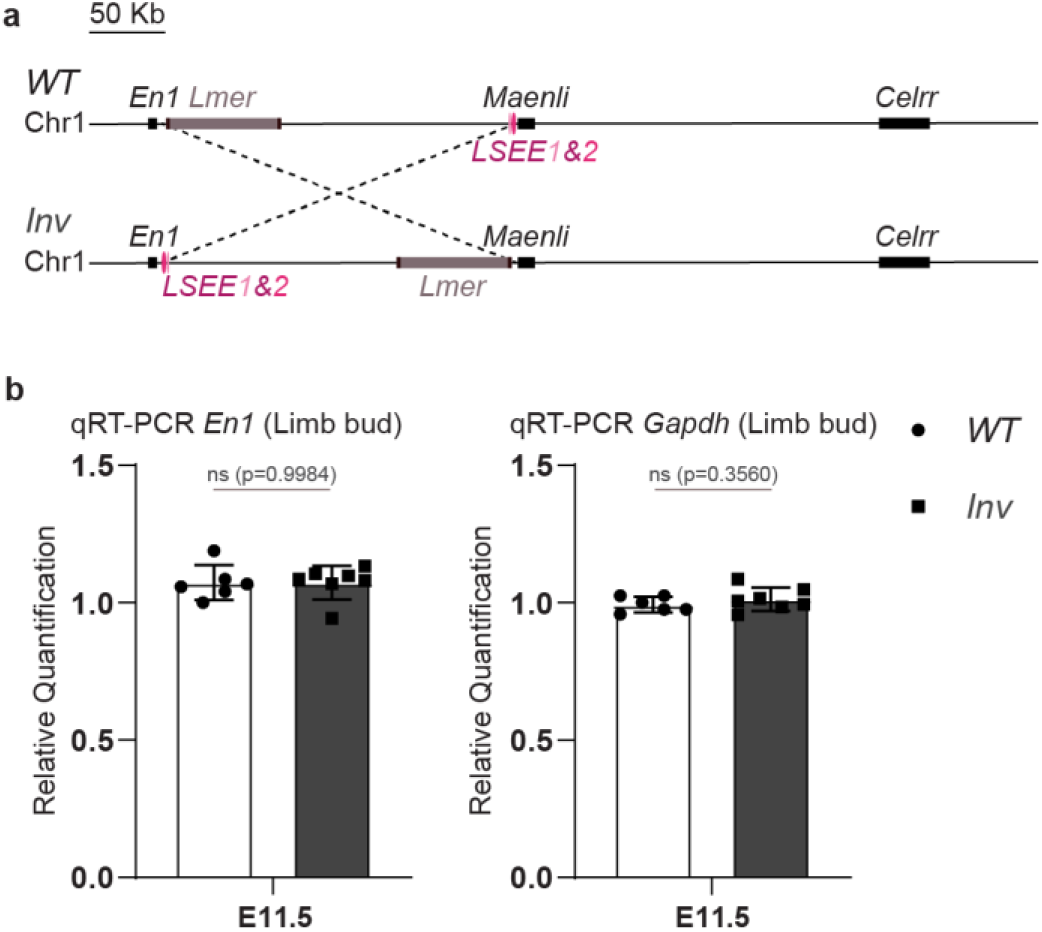
LSEE1&2 enhancer activity is not dependent on transcription elongation through the *Maenli* locus. **a**, Schematic representation of the CRISPR-Cas9 genetic inversion repositioning the LSEE1&2 enhancers away from *Maenli* within the *En1* regulatory landscape (*Inv*). **b,** Normalized qRT-PCRs of *En1* and *Gapdh* in E11.5 mouse limb embryos show no significant changes in *En1* and *Gapdh* expression upon repositioning the LSEE1&2 enhancers away from *Maenli* (*Inv*). Data were normalized to wild-type expression; one-tailed t-test; data are mean±SD; n= 6 *WT*, 7 *Inv* at E11.5 for *En1* and *Gapdh* expression quantification. ns, non-significant; p, p-value; WT, wild-type.

## Acknowledgments

Work in S.M. laboratory was supported by a grant from the Deutsche Forschungsgemeinschaft (DFG) (MU 880/16-1). Work in the laboratory of L.A. was supported by a UCL Award 156782/ Project 580721 and the Medical Research Council Laboratory of Medical Sciences which receives its core funding from the Medical Research Council. We are grateful to all members of the MPIMG transgene and mouse facility for embryonic stem cell aggregation and mouse husbandry and to all the MPIMG sequencing core facility members.

## Author Contributions

L.A. conceived the project. S.M. financed the majority of the project. A.R.R. and L.A. generated all the transgenic mouse models. L.W. performed morula aggregation. L.A. performed RNA-seq, qRT-PCRs, skeletal preparations, and phenotype analyses. A.M. performed ChIP-seq and ATAC-seq with the computational analyses performed by L.A. and A.R.R. L.A. performed cHi-C and R.S. performed the computational analysis. M.K., A.C.S., and U.F. provided technical support. A.R.R., A.M., N.B., and L.A. wrote the manuscript with input from R.S. and S.M., and approved by every co-author.

## Materials and Methods

### Generation of CRISPR/Cas9 genetic deletions and inversion

Genome editing experiments were performed as previously described^30^. sgRNAs were designed within close proximity of the deletion or inversion breakpoints, using the https://www.benchling.com/ platform to obtain candidate sgRNA sequences. To minimize off target effects, guide sequences were chosen to have a quality score above 95%. Complementary strands were annealed, phosphorylated, and cloned into the BbsI site of pX459 CRISPR/Cas vector (Addgene). The sequences of all sgRNAs used in this study are listed in Supplementary Table 1. G4 embryonic stem cells^31^ (ESCs) (400,000) (129/Sv X C57BL/6 F1 hybrid) were seeded on CD1 mouse embryonic fibroblast (MEF) feeders and cultured under standard ESC culture conditions. The cells were transfected with 8 µg of each pX459-sgRNA construct using FUGENE HD reagent (Promega) under manufacturer conditions. After 12 hours (h), cells were split, transferred into DR4 puromycin-resistant feeders, and selected with puromycin at a final concentration of 2 µg/ml for 48 h. Clones were then grown for 5 to 6 more days, picked, and transferred onto 96-well plates on CD1 feeders. After 48h of culture, plates were split in triplicates, two for freezing and one for growth and DNA harvesting. All clones were genotyped by PCR and quantitative PCR (qPCR) analyses. Positive clones were thawed and grown on CD1 feeders until they reached an average of four million cells. Three vials were frozen, and DNA was harvested from the rest of the cells to confirm the genotyping results.

ESCs and feeder cells were tested for *Mycoplasma* contamination using the MycoAlert detection kit (Lonza) and MycoAlert Assay Control Set (Lonza). DR4 and CD1 feeder cell lines were directly produced from mouse embryos originating from DR4 and CD1 mice mattings, respectively.

### Genotyping by polymerase chain reaction (PCR) and sequencing

Primers were designed at a distance of 400–500 bp from each cutting site, on both sides of sgRNA targets. Each allele has thus a set of four primers: F1(fwd)/R1(rev) amplifying one targeted site and F2(fwd)/R2(rev) amplifying the other targeted site. Deletions, duplications, and inversions were detected using the following set of primers: F1/R2, F2/R1, and F1/F2 and R1/R2, respectively. We genotyped each clone by running the PCR products on agarose gels and comparing PCR amplicon sizes. Clones showing evidence for the presence of the desired genomic rearrangement were further genotyped by Sanger sequencing. The sequences of all genotyping primers used in this study are listed in Supplementary Table 1.

### Genotyping by quantitative PCR (qPCR)

qPCR was performed on genomic DNA, using the QuantStudio 7 Flex Real-Time PCR System (v1.7.1) (Applied Biosystems). For the genotyping of copy number variations (CNVs), we designed different sets of primers. qPCR was carried out in a total volume of 20 µl containing 10 µl of Power Sybr Green Master Mix (Applied Biosystems), 0.4 µΜ of each primer and 10 ng of genomic DNA. Thermal cycling conditions were 95°C for 20 seconds (sec), followed by 40 cycles with 95°C for 3 sec and 60°C for 30 sec. A non-coding region was selected as the control amplicon. Validation experiments demonstrated that amplification efficiency of the control and all target amplicons were approximately equal. All samples were run in triplicate. The dosage of each amplicon relative to the control amplicon and normalized to wild-type DNA was determined using the 2^-ΔΔCT^ method.

### Transgenic mouse strains

Mice were generated from the corresponding ESC clone by diploid or tetraploid aggregation^32^, after thawing a frozen ESC vial seeded on CD1 feeders and grown for 2 days. Female mice of the CD1 strain were used as foster mothers.

All animal procedures were conducted as approved by the local authorities (LAGeSo Berlin) under the license numbers G0243/18 and G0176/19. Routine bedding, food, and water changes were performed. Mice were housed in a centrally controlled environment with a 12-h light/12-h dark cycle, temperature of 20–22.2 °C, and humidity of 30–50%. All animal experiments followed all relevant guidelines and regulations.

### RNA isolation and quantitative reverse transcription PCR (qRT–PCR)

To quantify RNA levels in wild-type and mutant mice at E9.5, E10.5, and E11.5 developmental stages (G4 background), limb buds were micro-dissected in cold PBS, immediately snap-frozen, and stored at −80 °C. Total RNAs were extracted using the RNeasy Mini Kit (QIAGEN) according to the manufacturer’s instructions. RNAs were treated with DNase I (Thermo Fisher Scientific) at 37 °C for 15 min followed by 10 min incubation at 70 °C for DNase I inactivation. Complementary DNAs (cDNAs) were generated using the Superscript III First-Strand Synthesis System (Thermo Fisher Scientific) whereby 1 µg of RNA was reverse-transcribed using oligo(dT)20. To quantify the relative abundance of transcripts, qRT–PCR analyses were done in technical triplicates using the Power Sybr Green Master Mix (Applied Biosystems) as described above. The dosage of each amplicon was normalized to the endogenous control amplicon (*Rps9* cDNAs). The 2^-ΔΔ*CT*^ method^33^ was used to calculate the fold change between wild-type and mutant samples. Using a one-tailed Student’s *t*-test, we tested the ability of mutations to result in changes in *Gapdh*, *Maenli* and/or *En1* gene expression. The variance in mutant and wild-type samples was assumed equal. The sequences of all qRT-PCR primers used in this study are listed in Supplementary Table 1.

### Skeletal preparation

E18.5 fetuses (G4 background) were kept in H2O for 1–2 h at RT and heat-shocked at 65 °C for 1 min. The skin was taken off and the abdominal and thoracic viscera were removed using forceps. The fetuses were then fixed in 100% ethanol overnight. Afterwards, the cartilage was stained overnight using alcian blue staining solution (150 mg/l alcian blue 8GX in 80% ethanol and 20% acetic acid). Fetuses were then rinsed and post-fixed in 100% ethanol overnight. After 24 h, initial clearing was done by incubating the fetuses for 20 min in 1% potassium hydroxide in H2O, followed by alizarin red (50 mg/l alizarin red S in 0.2% potassium hydroxide) staining of bones overnight. Following this, rinsing and clearing was done for several days using low concentrations of potassium hydroxide. The stained embryos were dissected in 80% glycerol and limbs were imaged using a ZEISS SteREO Discovery.V12 with cold light source CL9000 microscope and Leica DFC420 digital camera.

### Phenotypic evaluation

Phenotypic analyses for 7-8 weeks mutant mouse lines were carried out for at least 10 animals per analysis.

### SureSelect design

The capture Hi-C SureSelect library probes were designed over the genomic interval (mm9, chr1: 119,650,000–124,400,000) using the SureDesign online tool from Agilent.

### Capture Hi-C (cHi-C)

cHi-C libraries were prepared from E10.5 wild-type limb buds (CD1 background). cHi-C experiments were performed as duplicates. Per biological replicate, 10 pairs E10.5 limb buds (2∼3 × 10^6^ cells) were micro-dissected in PBS at RT. A single-cell suspension was obtained by incubating the tissue for 10 min at 37 °C in 1 ml Gibco trypsin-EDTA 0.05% (Thermo Fisher Scientific). Cells were resuspended in 5 ml 10% fetal bovine serum/PBS and fixed by adding 5 ml 4% formaldehyde (Sigma-Aldrich) at a final concentration of 2%. Cells were mixed for 10 min at RT. Fixation was quenched using 1.425 M glycine (Merck) on ice and immediately centrifuged at 2100 r.p.m. for 8 min. Supernatant was removed and the pellet resuspended in lysis buffer (final concentration of 10 mM Tris, pH 7.5, 10 mM NaCl, 5 mM MgCl2, 0.1 M EDTA, and 1× cOmplete protease inhibitors (Sigma-Aldrich)) and incubated on ice for 10 min. Cells were then centrifuged at 2900 r.p.m. for 5 min at 4 °C, followed by removal of supernatant, snap-freezing, and storage at −80 °C. For the preparation of the 3C library, the pellet was resuspended in 60 µl 10× DpnII buffer (Thermo Fisher Scientific), and incubated with 15 µl 10% SDS for 1 h at 37 °C and 900 r.p.m. 150 µl 10% Triton X-100 was then added and the pellet was incubated for 1 h at 37 °C and 900 r.p.m. 600 µl of 1× DpnII buffer was added to the samples. A 10-µl aliquot was taken as undigested control and stored at −20 °C. The chromatin was digested using 40 µl 10 U/µl DpnII for 4 h at 37 °C and 900 r.p.m.; another 20 µl of DpnII enzyme was then added and samples were incubated overnight at 37 °C with rotation. After overnight incubation, samples were supplemented with 20 µl DpnII enzyme and incubated for four more hours at 37 °C with rotation. The DpnII restriction enzyme was inactivated at 65 °C for 20 min. Next, the digested chromatin was diluted and religated in 5.1 ml H2O, 700 µl 10× ligation buffer (Thermo Fisher Scientific), and 1.67 µl 30 U/µl T4 DNA ligase (Thermo Fisher Scientific). Ligation reactions were incubated at 16 °C overnight with rotation. A 100-µl aliquot was taken to test ligation efficiency and stored at −20 °C. The chimeric chromatin products and test aliquots were de-cross-linked overnight by adding 30 µl and 5 µl 20 mg/ml proteinase K, respectively, and incubated at 65 °C overnight. Following this, 30 µl or 5 µl 10 mg/ml RNase A was added to the samples and aliquots, respectively, and incubated for 45 min at 37 °C. Chromatin was then precipitated by adding 1 volume phenol-chloroform to the samples and aliquots, vigorously shaking them, followed by centrifugation at 3,750 r.p.m. at RT for 15 min. The upper phase containing the chromatin was transferred to a new tube and the volume was adjusted to 7 ml with H2O. Samples were then supplemented with 1 ml 3M NaAc, pH 5.6, 35 ml 100% ethanol, and 7 µl 20 mg/ml glycogen and frozen overnight at −80 °C. The precipitated chromatin was isolated by centrifugation at 8,350 g for 20 min at 4 °C. The chromatin pellet was washed with 10 ml 70% ethanol and further centrifuged at 3,300 g for 15 min at 4 °C. Finally, the 3C library chromatin pellet was dried at RT and resuspended in 150 µl 10 mM Tris-HCl, pH 7.5 at 37 °C. To check the 3C library, the undigested, digested, and ligated aliquots were loaded on a 1% agarose gel. The 3C library was then sheared using a Covaris sonicator (duty cycle: 10%; intensity: 5; cycles per burst: 200; time: 6 cycles of 60 s each; set mode: frequency sweeping; temperature: 4-7 °C). Adaptors were added to the sheared DNA and amplified according to the manufacturer’s instructions for Illumina sequencing (Agilent). The library was hybridized to the custom-designed SureSelect beads and indexed for sequencing (150 bp paired-end) following the manufacturer’s instructions (Agilent).

### cHi-C processing

FASTQ files were processed with the HiCUP pipeline v0.6.1^34^ (no size selection, Nofill: 1, Format: Sanger) using Bowtie2 v2.3.4.1^35^ for mapping short reads to the reference genome mm9. After the mapping, filtering and deduplication steps of the HiCUP pipeline, replicates were merged by combining their bam files. Juicer tools v1.7.6^36^ was used to generate binned contact maps and to normalize maps by Knights and Ruiz (KR) matrix balancing^37^. For the generation of the contact maps, only the genomic region (chr1:120,200,001-124,400,000, mm9), part of the region enriched in the capturing step, was considered. Therefore, only read-pairs mapping to the region of interest were kept and their coordinates were shifted towards the origin by the offset of this region. Afterwards, cHi-C maps were generated with the Juicer tools ‘pre’ command using a custom chrom.sizes file containing only the length of the genomic region of interest (4.2 Mb) and applying a minimum MAPQ of 30. After exporting KR normalized cHi-C maps at 10kb bin size, coordinates were shifted back to mm9 coordinates. cHi-C maps were displayed as heatmaps in which values above the 98.5^th^ percentile were truncated to improve visualization.

### Virtual 4C profile generation

In order to obtain more fine-grained interaction profile for the *En1* promoter, a virtual 4C-like profile was generated from mapped, filtered and deduplicated cHi-C read pairs obtained from the HiCUP pipeline. A read-pair was considered in the profile, when it had a MAPQ≥30 and one read mapped to the defined viewpoint region while the other one mapped outside of it. Out of these pairs, reads mapping outside of the viewpoint region were counted per restriction fragment and binned to a 1kb grid. The count value of restriction fragments spanning more than one bin was distributed proportionally to the corresponding bins. Afterwards, the profile was smoothed by averaging over five bins and scaled by the factor 10^3^ / counts within the enriched region. For the computation of the scaling factor, the viewpoint region and a margin ±5 kb around it were excluded. The processing was performed with custom Java code using htsjdk v.2.12.0 (https://samtools.github.io/htsjdk/).

### RNA-sequencing (RNA-seq)

E9.5 forelimb buds, and E10.5 and E11.5 limb buds were micro-dissected from wild-type and mutant embryos (G4 background) in cold PBS, immediately snap-frozen, and stored at –80 °C. RNA-seq experiments were performed in duplicates. Total RNAs were extracted using the RNeasy Mini Kit (QIAGEN) according to the manufacturer’s instructions. RNAs were treated with DNase I (Thermo Fisher Scientific) at 37 °C for 15 min followed by 10 min incubation at 70 °C for DNase I inactivation. Samples were poly-A enriched and sequenced (150 pb paired-end) using Illumina technology following standard protocols.

### RNA-seq data processing

Paired-end RNA-seq reads were mapped to a custom mouse reference genome (mm9), including *Maenli* coordinates (chr1:122,744,478-122,755,676) using the STAR mapper^38^ (version 2.6.1d) (splice junctions based on RefSeq; options: –-alignIntronMin 20 – alignIntronMax 500000 – outFilterMultimapNmax 5 – outFilterMismatchNmax 10 –outFilterMismatchNoverLmax 0.1).

### ATAC-seq data processing

Raw sequencing files were processed with Cutadapt to trim the adapter sequence^39^, Bowtie2 was used for mapping^39^. SAMtools was used for filtering, sorting, and duplicate removal^40^. deepTools was used to create coverage tracks^41^. Calling of peaks was performed using Genrich with default settings (https://github.com/jsh58/Genrich)^42^.

### Data Availability Statement

All datasets have been deposited in the Gene Expression Omnibus (GEO) database and are accessible under GSE277393, GSE277394, and GSE277395 (or published previously under GSE137335 and GSE84795).

**Supplementary Table 1.**
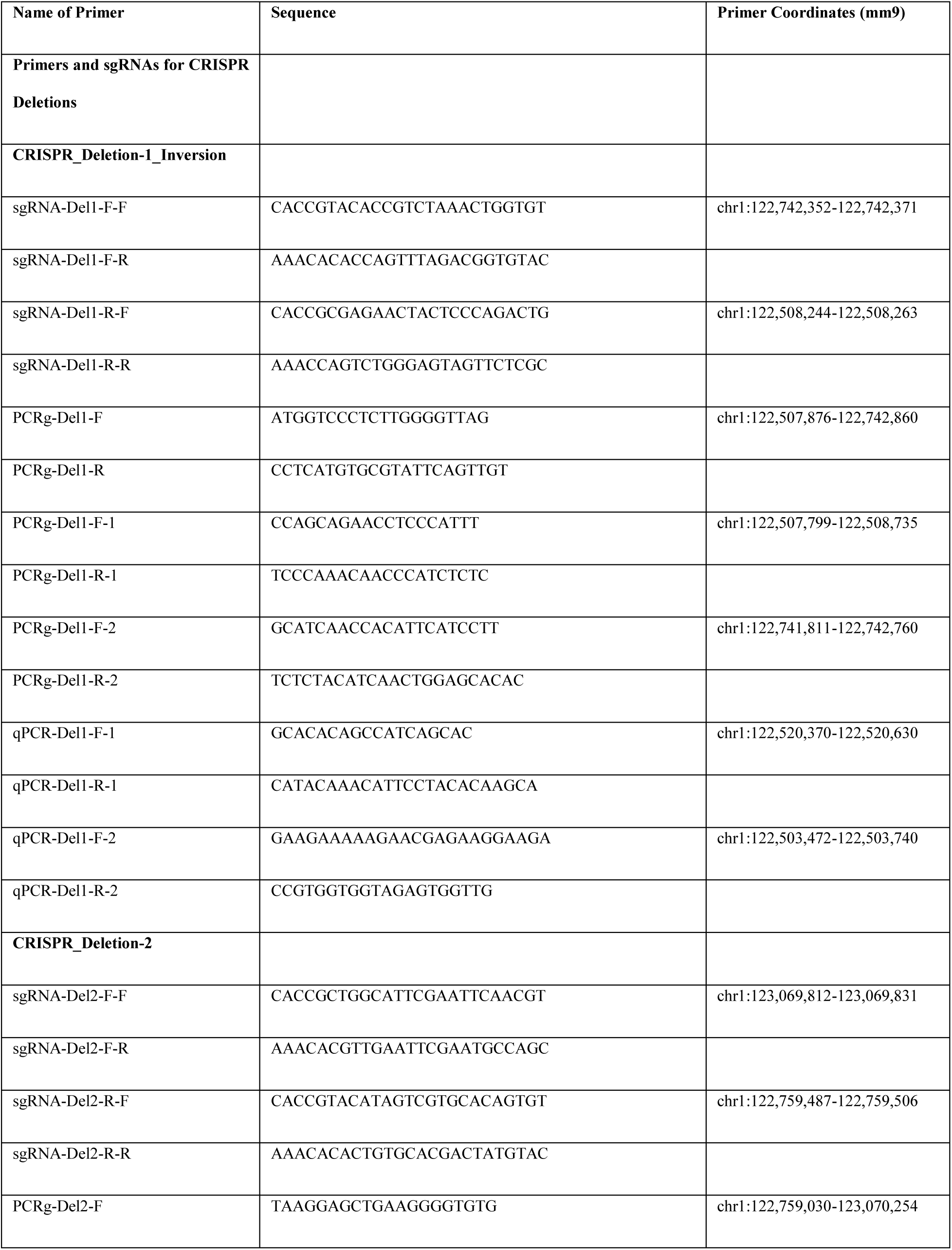
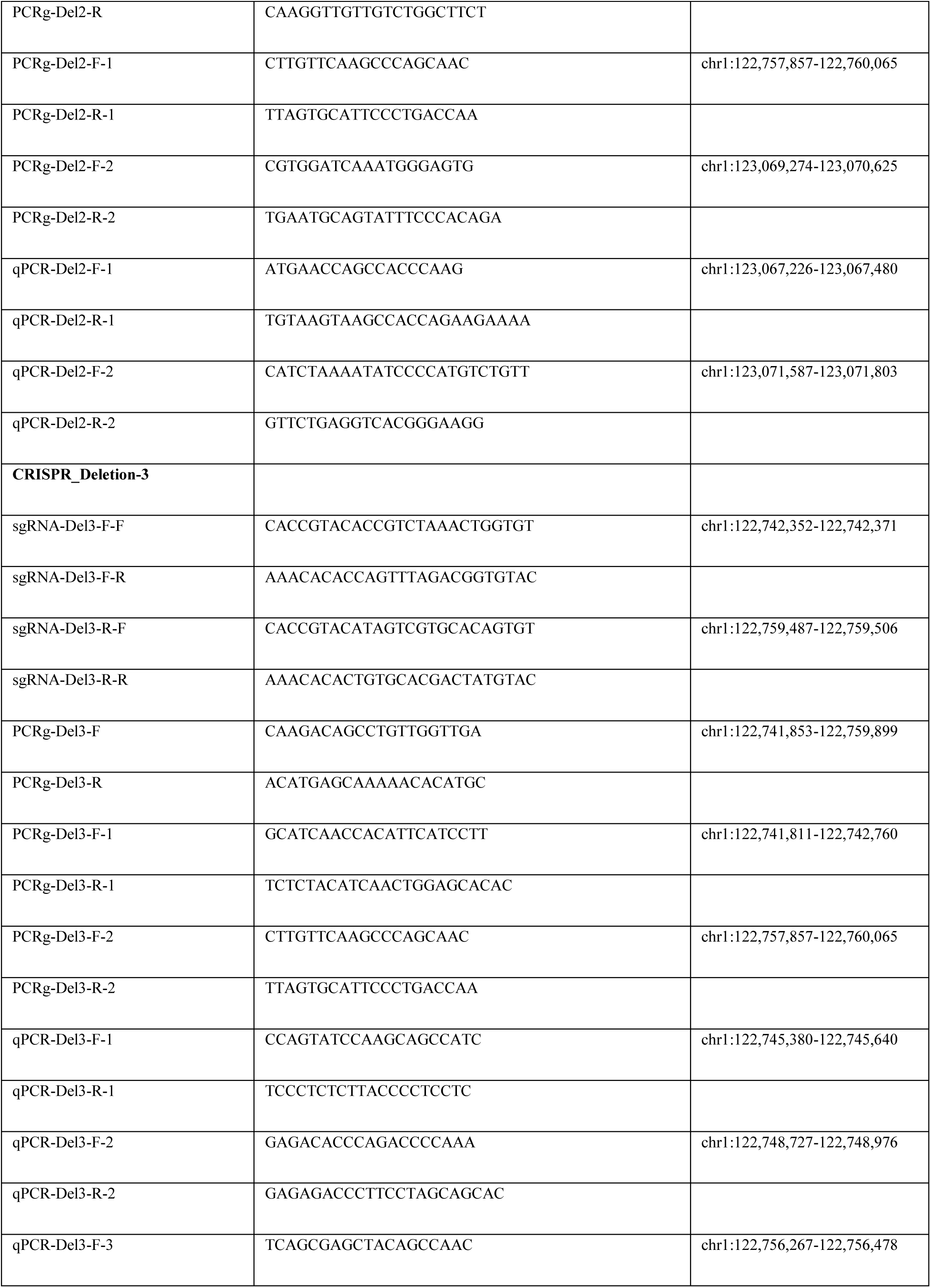
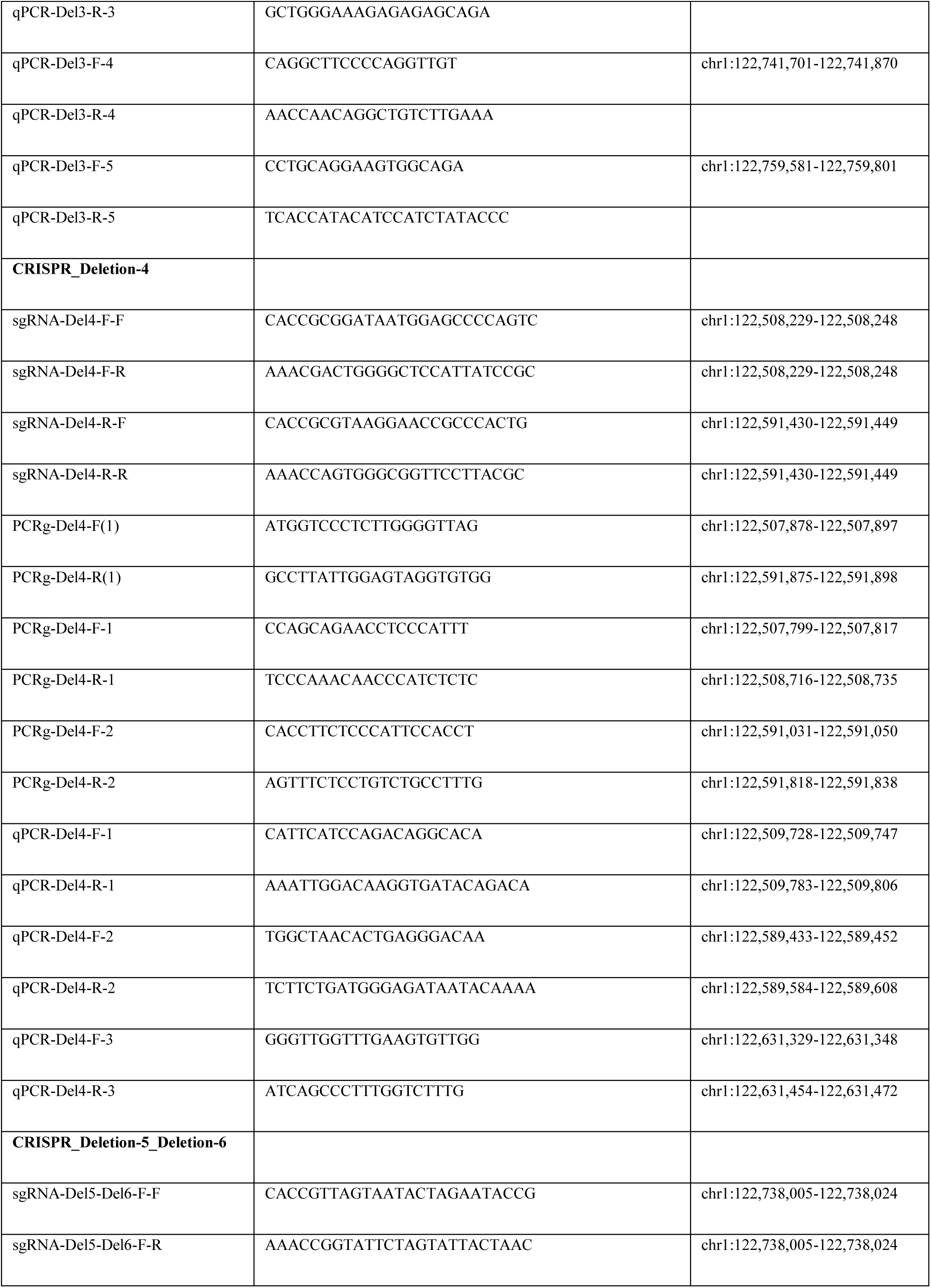
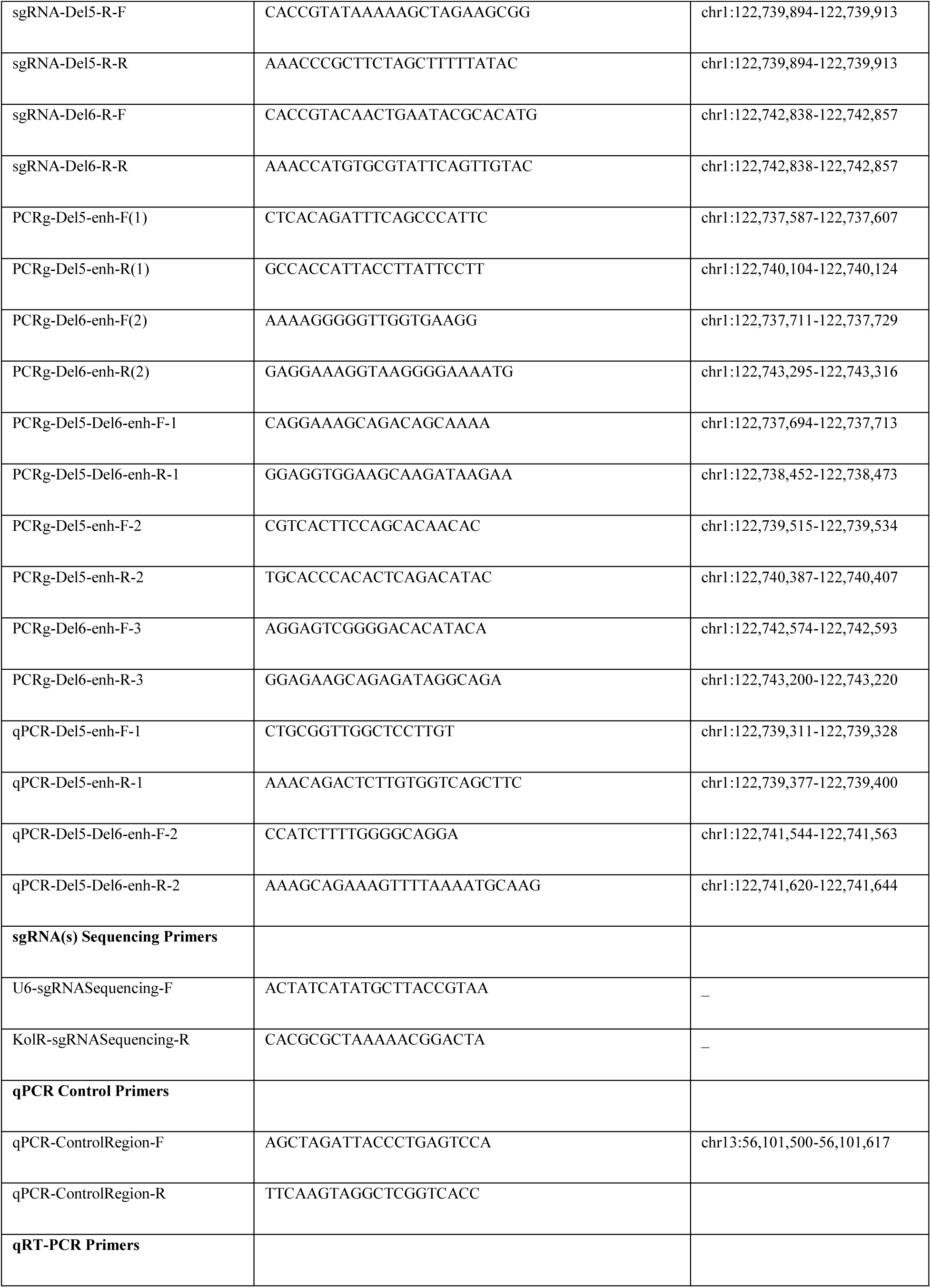
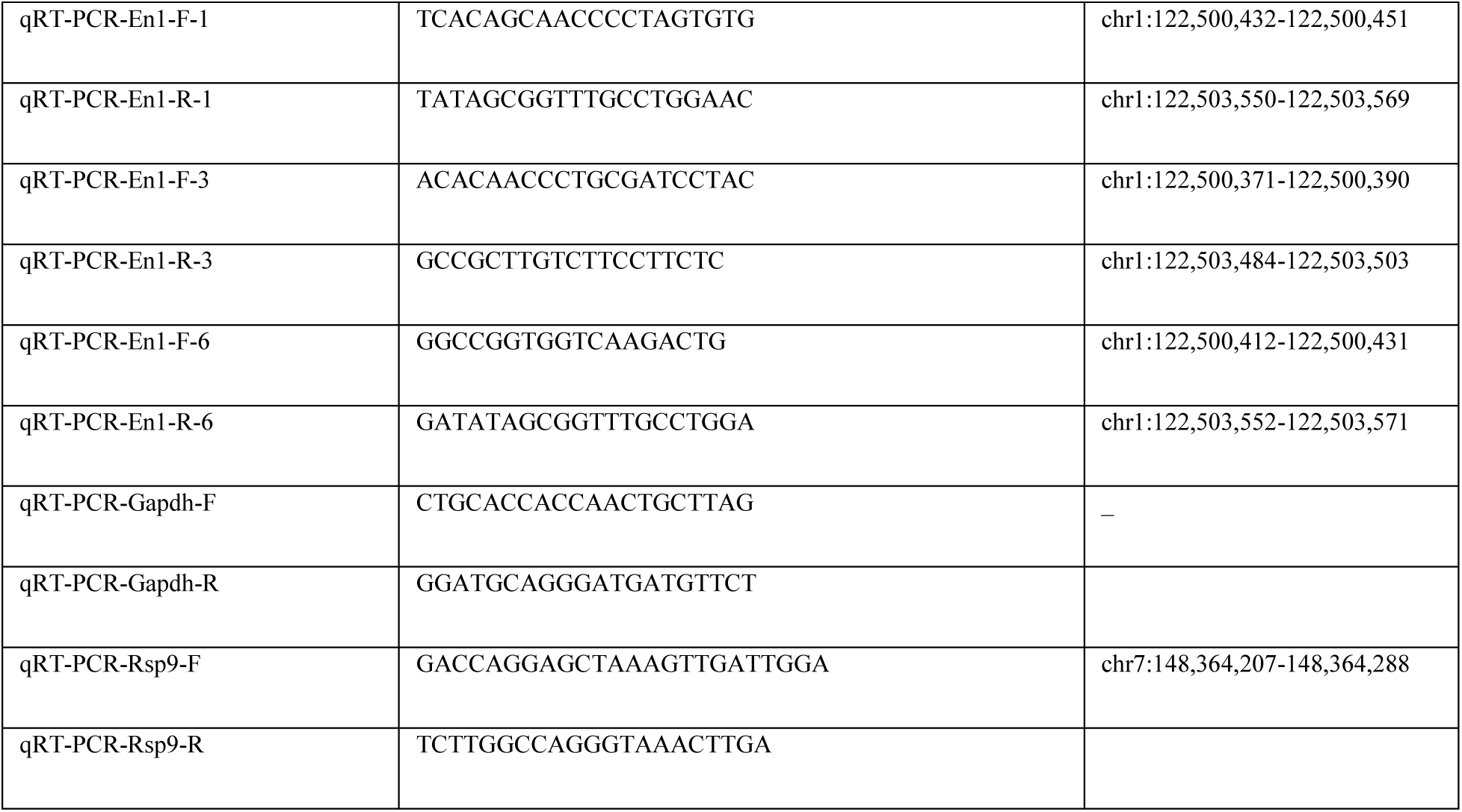
List of primer sequences and sgRNAs for CRISPR targeting.

